# Immune modulation to improve survival of respiratory virus infections in mice

**DOI:** 10.1101/2020.04.16.045054

**Authors:** Shradha Wali, Jose R. Flores, Ana Maria Jaramillo, David L. Goldblatt, Jezreel Pantaleón García, Michael J. Tuvim, Burton F. Dickey, Scott E. Evans

**Affiliations:** University of Texas MD Anderson Cancer Center UTHealth Graduate School of Biomedical Sciences, Houston Texas 77030; University of Texas MD Anderson Cancer Center, Department of Pulmonary Medicine, Houston Texas 77030

**Keywords:** Immunomodulation, immunopathology, CD8^+^ T cells, viral pneumonia, inducible epithelial resistance

## Abstract

Viral pneumonia remains a global health threat requiring novel treatment strategies, as strikingly exemplified in the SARS-CoV-2 pandemic of 2019-2020. We have reported that mice treated with a combination of inhaled Toll-like receptor (TLR) 2/6 and TLR 9 agonists (Pam2-ODN) to stimulate innate immunity are broadly protected against respiratory pathogens, but the mechanisms underlying this protection remain incompletely elucidated. Here, we show in a lethal paramyxovirus model that Pam2-ODN-enhanced survival is associated with robust virus inactivation by reactive oxygen species (ROS), which occurs prior to internalization by lung epithelial cells. However, we also found that mortality in sham-treated mice temporally corresponded with CD8^+^ T cell-enriched lung inflammation that peaks on days 11-12 after viral challenge, when the viral burden has waned to a scarcely detectable level. Pam2-ODN treatment blocked this injurious inflammation by reducing the viral burden, and alternatively, depleting CD8^+^ T cells 8 days after viral challenge also decreased mortality. These findings reveal opportunities for targeted immunomodulation to protect susceptible individuals against the morbidity and mortality of respiratory viral infections.

## Introduction

Viruses are the most frequent cause of community acquired pneumonia in children and adults, resulting in significant morbidity in vulnerable subjects and exerting a tremendous health care burden (1–5). Seasonal influenza and emergent pandemic viruses, such as SARS-CoV-2, inflict particular mortality in susceptible individuals, with clinicians frequently lacking effective interventions to improve patient outcomes (6–9). Moreover, in addition to causing acute disease, respiratory virus infections are often complicated by chronic lung pathologies, such as asthma induction, progression and exacerbation (10–12). Therefore, development of novel therapeutic anti-viral strategies is required to effectively prevent and treat respiratory infections and their associated chronic complications (13, 14).

While lung epithelial cells are the principal targets of most respiratory viruses (15), there is expanding evidence that lung epithelia themselves are capable of generating anti-microbial responses (12, 16, 17). We hypothesized that lung epithelial cells can be harnessed to control virus replication, thereby enhancing acute survival and reducing chronic complications of virus infections (18–21). Our group has previously described the phenomenon of inducible epithelial resistance wherein the lungs’ mucosal defenses can be broadly stimulated to protect against a wide range of respiratory pathogens, including viruses (18–23). This protection is induced by a single inhalation of a combination treatment consisting of Toll like receptor (TLR) 2/6 and 9 agonists (Pam2-ODN) shortly before or after viral challenge. While no individual leukocyte populations have been identified as critical for Pam2-ODN-induced resistance, lung epithelial cells are essential to the inducible anti-viral response (18). Further, we have shown that Pam2-ODN mediated protection is dependent upon epithelial generation of reactive oxygen species (ROS) but, interestingly, does not require Type I interferons (22, 23). More recently, we have shown prevention of chronic virus-induced asthma in mice treated with Pam2-ODN but we have not clarified the anti-viral mechanisms (24).

In this study, we investigated the mechanisms of Pam2-ODN enhanced mouse survival of pneumonia caused by a paramyxovirus, Sendai virus (SeV). We found that Pam2-ODN treatment not only reduced lung SeV burden but also decreased epithelial cell injury and host immunopathologic leukocyte responses to SeV infections. While CD8^+^ T cells are known to contribute to anti-viral immunity, it is shown here that CD8^+^ T cells contribute substantially to mortality, and this effect can be prevented by Pam2-ODN treatment early in the course of infection or CD8^+^ depletion late in the course. Further, we demonstrate anti-viral mechanisms of inducible epithelial resistance, where virus particles are inactivated in a ROS-dependent manner prior to internalization by their epithelial targets.

## Materials and Methods

### Mice

All *in vivo* experiments were performed using 6- to 10-week-old C57BL/6J mice of a single sex and were handled according to the Institutional Animal Care and Use Committee of MD Anderson Cancer Center, protocol 00000907-RN01.

### Cells

Mouse lung epithelial (MLE-15) cells were kindly provided by Jeffrey Whitsett, Cincinnati Children’s Hospital Medical Center. Mouse tracheal epithelial cells were harvested and cultured as previously described (22, 25). See *Supplemental Methods* for additional details.

### TLR treatments and viral challenges

Cells were treated with Pam2CSK_4_ (2.2 μM) and ODN M362 (0.55 μM) as previously described (22, 23). Mice were treated with 10 ml of Pam2CSK_4_ (4 μM) and ODN M362 (1 μM) by nebulization as previously described (22, 23). For *in vitro* challenges, SeV at multiplicity of infection (MOI) = 1 was used. Unless otherwise stated, mice were challenged with 1 × 10^8^ plaque forming units (pfu) inserted into the oropharynx as described (24). See *Supplemental Methods* for additional details.

### Flow cytometry

Single cells from disaggregated lungs or cell culture were stained as indicated in the antibody table (Table 1), fixed, and acquired on a BD LSRII (BD Biosciences). See *Supplemental Methods* for additional details.

**Table 1.**

| **Antibodies** | **Vendor** | **Catalogue numbers** |
| --- | --- | --- |
| CD3 | Tonbo | 65-0031-U100 |
| CD4 | Tonbo | 60-0042-U100 |
| CD8 | Tonbo | 25-0081-U100 |
| Live dead | Tonbo | 13-0870-T500 |
| CD25 | Biolegend | 102038 |
| Foxp3 Treg kit | eBiosciences | 72-5775 |
| CD8-Depleting Ab | Bioxell | BE0223-A025 |
| CD19 | Biolegend | 115507 |
| B220 | BD Biosciences | 562922 |
| Anti-SeV virus Ab | MBL International | PD029 |
| Ki67 | Invitrogen | MA5-14520 |
| cCasp3 | Cell signaling | 9662S |

### Epithelial proliferation assays

Epithelial proliferation was determined by staining lung sections for EdU 24 h after intraperitoneal injection. See Supplemental *Methods* for details.

### CD8^+^ T cell depletion

Anti-CD8-β antibody (200 μg/mouse, clone 53-5.8, Bioxell) was delivered to mice intraperitoneally at indicated time points. CD8^+^ T cell depletion was confirmed by flow cytometry analysis 24 to 48 h after depletion.

### Viral burden quantification

Viral burden was determined by reverse transcription quantitative PCR (RT-qPCR) of the Sendai Matrix (M) protein normalized to host house-keeping gene 18SRNA. For *in vivo* experiments, mouse lungs were collected 5 days after SeV challenge. For in vitro experiments, cell lysates were collected 24 h after infection, unless otherwise indicated. See *Supplemental Methods* for additional details.

### ROS inhibition *in vitro* and *in vivo*

NADPH oxidase activity was inhibited using GKT137831 (Selleckchem). Mitochondrial ROS production was inhibited using the combination of FCCP (Cayman Chemicals) and TTFA (Cayman Chemicals). See *Supplemental Methods* for additional details.

### Viral attachment assays

For most enveloped viruses, internalization into epithelial cells is inhibited at 4° C without affecting viral binding to epithelial cells (26–28). MLE-15 cells were infected with SeV at 4° C for 4 h, washed to remove unattached virus, then assessed for uninternalized SeV burden using immunofluorescence or flow cytometry. See *Supplemental Methods* for additional details.

## Results

### Enhanced mouse survival of SeV infection by Pam2-ODN treatment

Aerosolized Pam2-ODN treatment one day prior to SeV challenge increased mouse survival of SeV challenge (Figure 1A), similar to the protection observed against lethal influenza pneumonia (18, 21, 22). The survival benefit was associated with reduced lung SeV burden, as measured by SeV M gene expression (Figure 1B). Investigating the natural progression of infection revealed that SeV lung burden was maximal on day 5 and gradually decreased until falling below the limit of quantification (LOQ) by day 11 (Figure 1C). Pam2-ODN pretreatment reduced SeV burden on all assessed days (Figure 1C). Although the lethality of SeV infection was exquisitely dependent on the inoculum size, we strikingly found that peak mortality paradoxically occurred around days 10 to 12 after infection irrespective of inoculum size, despite the fact that SeV is essentially undetectable that long after challenge (Figure 1A, C, and D).

**Figure 1.**
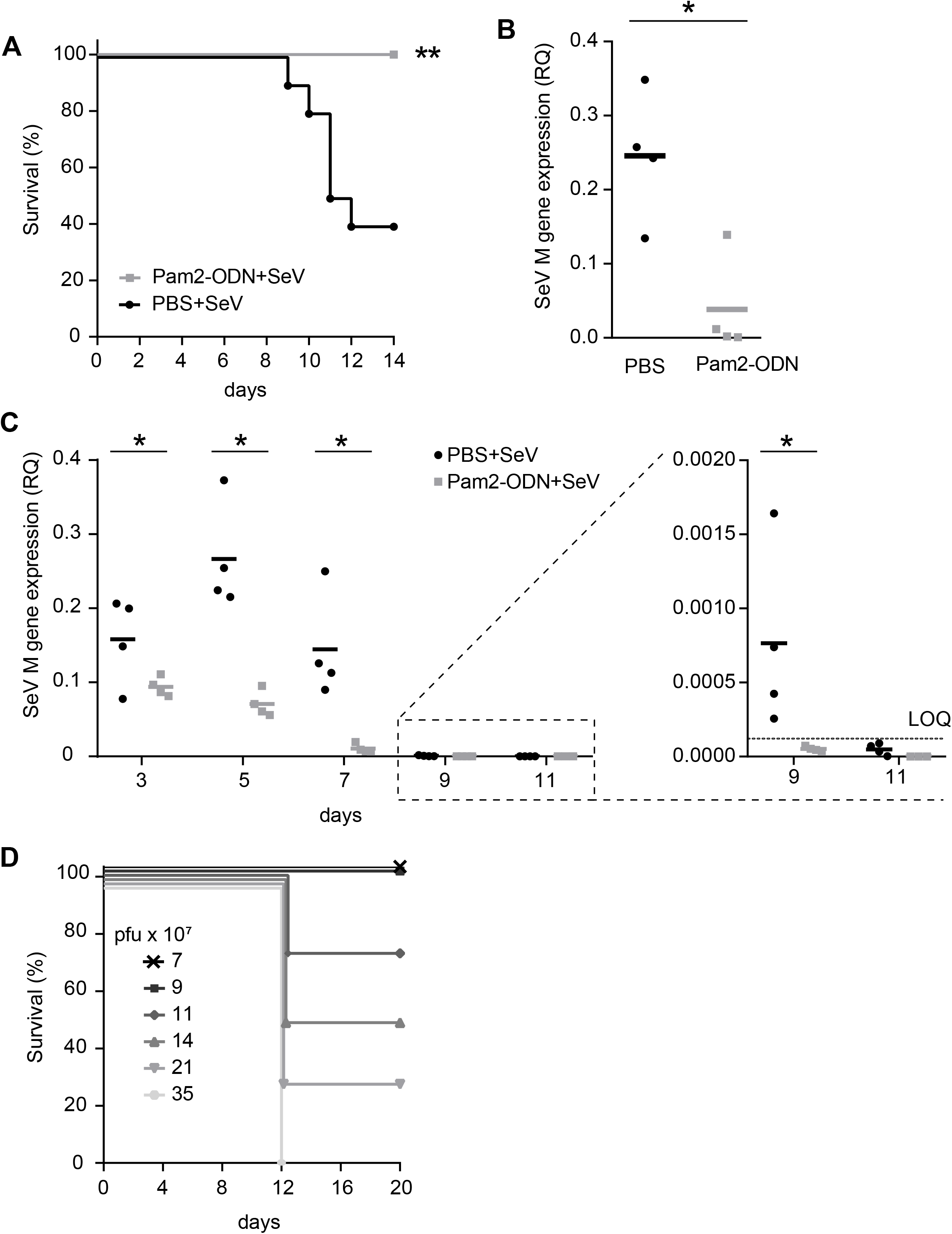
Pam2-ODN enhances mouse survival of SeV infection and reduces lung virus burden. **(A)** Survival of mice treated with PBS or Pam2-ODN one day prior to SeV virus challenge. **(B)** Mouse lung SeV burden 5 days after infection assessed by qPCR for Sendai Matrix (M) gene (Relative quantification, RQ to 18S) relative to 18S. **(C)** Time course of lung SeV burden in mice treated with PBS or Pam2-ODN. **(D)** SeV inoculum dependent mouse survival. Data are representative from three independent experiments. n=10 mice per group in survival plots, n=4 mice/group in virus burden experiments. LOQ, limit of quantification. \**p*<0.05, \*\**p*<0.005.

### Pam2-ODN treatment attenuates SeV-induced epithelial injury

This temporal dissociation between peak virus burden and peak mortality led to the hypothesis that SeV-induced mortality may not be exclusively driven by excessive virus burden but may also result from untoward SeV-induced host immune response. Therefore, the acute changes in mouse lungs following SeV infection were characterized. We found increases in lung epithelial cleaved caspase 3 (cCasp3), a marker for programmed cell death, on days 7 to 11 after SeV infection (Figure 2A, upper panel). Virus infection-related epithelial cell injury and death is typically associated with proliferative repair mechanisms (29, 30). Staining the infected mouse lung tissue for Ki67 and EdU revealed maximum signals for both markers in the second week after infection (Figure 2B-E, upper panel). These events of lung epithelial cell death and proliferation coincided with the peak of mortality (day 12, Figure 1E). Further, hematoxylin and eosin staining of lung tissues infected with SeV showed profound increases in inflammatory cells from days 7 to 10 with evidence of damaged airway and parenchymal tissue (Figure 2F). However, Pam2-ODN pretreatment of mice reduced epithelial cell injury and proliferation (Figure 2A-E, lower panel). This temporal association of epithelial injury and death after viral clearance supported our hypothesis that mouse mortality caused by SeV infection is due in part to the host immune response to SeV infections.

**Figure 2.**
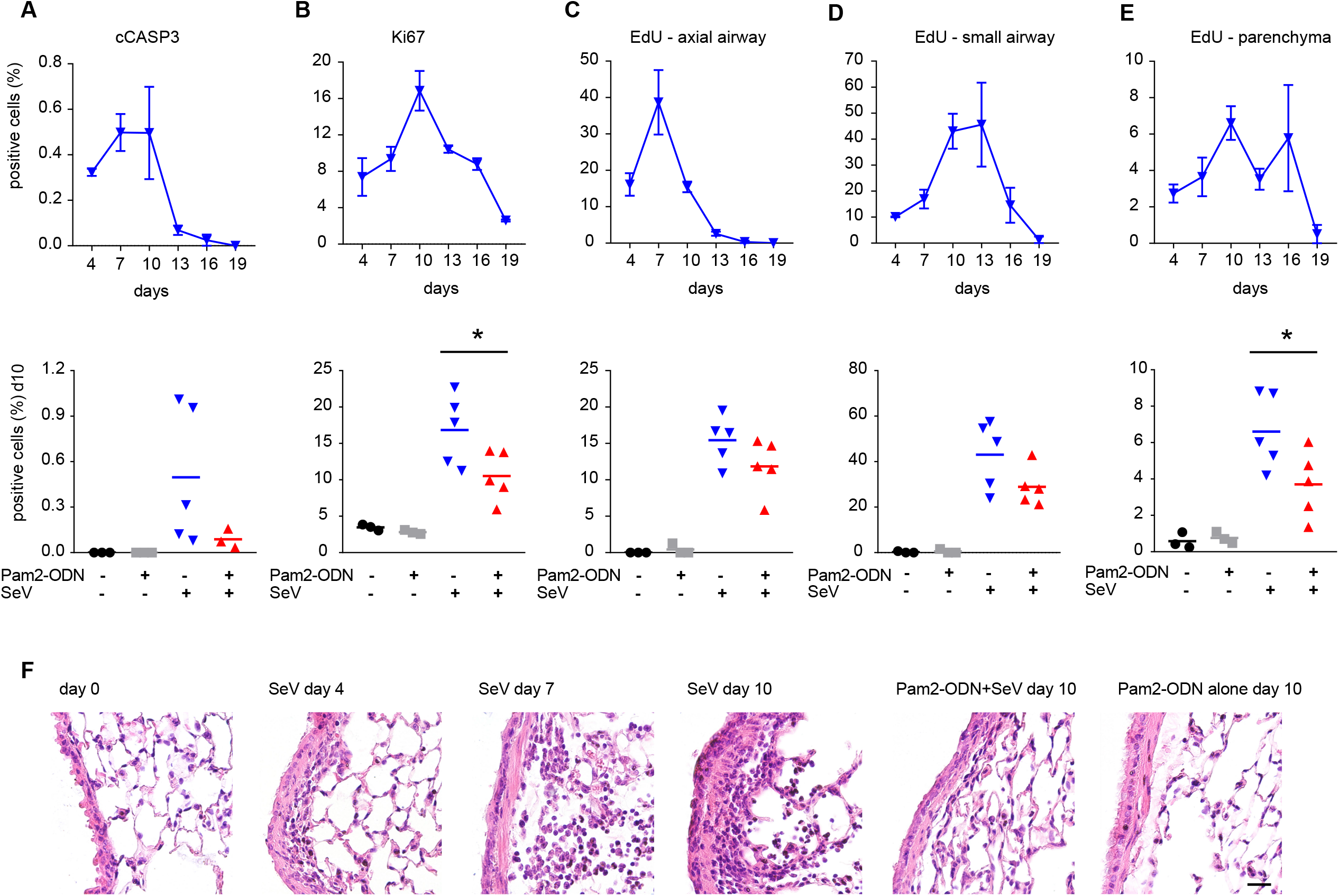
Pam2-ODN pretreatment reduces epithelial cell death and proliferation during acute SeV infection. Cleaved caspase 3 (cCasp3) **(A)** or Ki67 **(B)** positive cells in mouse lung epithelium after SeV infection with or without Pam2-ODN treatment (lower panel). EdU positive cells in axial **(C)**, small airways **(D)** and parenchyma **(E)** after SeV infection with or without Pam2-ODN (lower panel). **(F)** Mouse lung histology following SeV challenge with or without Pam2-ODN. n=5 mice per condition. Data are representative from two independent experiments. Scale bar = 100 μm. \**p*<0.05.

### Pam2-ODN attenuates SeV-induced lymphocytic lung inflammation

To explore this hypothesis, the host leukocyte response to SeV infection was characterized. Differential Giemsa staining of bronchoalveolar lavage (BAL) cells revealed increased neutrophils on days 2 to 5 and increased macrophages on days 5 to 8 (Figure 3A, left and middle panel, solid grey line) after SeV challenge. Congruent with our prior studies, inhaled treatment with Pam2-ODN in the absence of infection led to a rapid rise in neutrophils that was resolved within 5 days (Figure 3A, dashed line) (31). The neutrophil response to SeV challenge was modestly increased among mice pretreated with Pam2-ODN (Figure 3A, left panel, solid dark line). Pam2-ODN-treated, SeV-challenged mice showed almost no difference in macrophage number compared to PBS-treated, SeV-challenged mice (Figure 3A, middle panel, solid dark line). A rise in lymphocytes was observed on days 8 to 11 in PBS-treated, SeV-challenged mice (Figure 3A, right panel, solid grey line), temporally corresponding with peak mortality. However, Pam2-ODN treated, SeV-challenged mice displayed significantly reduced lymphocyte numbers at every time point assessed (Figure 3A, right panel, solid dark line). The gating strategy for lymphocyte subsets by flow cytometry is shown in Supplementary Figure 1. A modest reduction in CD4^+^ T cells was observed in Pam2-ODN-treated, SeV-challenged mice compared to PBS-treated, SeV-challenged mice (Supplementary Figure 2). We also found the percentage of CD19^+^ B220^+^ B cells reduced after SeV infection in comparison to Pam2-ODN treated and uninfected mice (Supplementary Figure 2), as has been seen with other viral models (32, 33). However, the biggest difference between groups was in CD8^+^ T cells, with Pam2-ODN-treated, SeV-challenged mice displaying a significantly lower number and percentage of CD8^+^ T cells than PBS treated, SeV-challenged mice (Figure 3B, C). Since the greatest difference after Pam2-ODN treatment was in CD8^+^ T cell levels and there was a tight correlation between peak mortality and the increase in lung CD8^+^ T cells on days 8 to 11, we investigated the role of CD8^+^ T cells in SeV-induced mortality.

**Figure 3.**
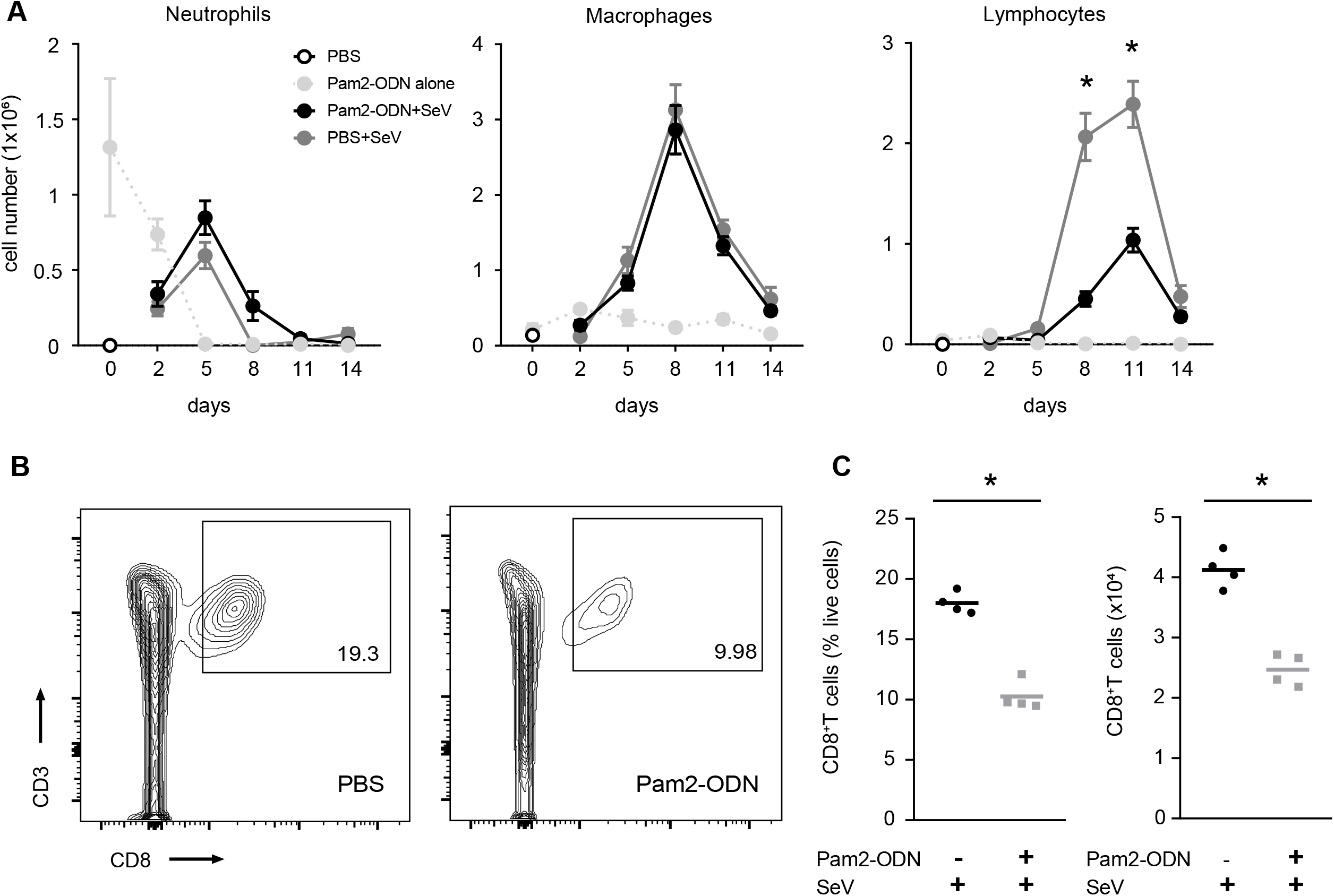
Pam2-ODN pretreatment reduces SeV induced lung CD8^+^ T cells. **(A)** Differential Giemsa staining of BAL cells from mice challenged with SeV with or without Pam2-ODN pretreatment. **(B)** Flow cytometry for CD8^+^ T cells from disaggregated mouse lungs 11 days after SeV infection with or without Pam2-ODN. **(C)** Lung CD8^+^ T cells 11 days after SeV challenge in mice pretreated with PBS or Pam2-ODN. Data are representative of three independent experiments for (A) and of five independent experiments for (B) and (C). \**p*<0.05 compared to PBS+SeV

### Depleting CD8^+^ T cells after viral clearance enhances survival of SeV infection

To understand the apparent contributions of host immunopathology to mouse outcomes, we depleted CD8^+^ T cells on day 8 -- after virus burden was substantially reduced but before peak mouse mortality (Figures 1 and 4A). Mice depleted of CD8^+^ T cells displayed significantly enhanced survival of SeV challenge compared to mice with intact CD8^+^ T cells (Figure 4B). Depletion of CD8^+^ T cells was confirmed by flow cytometry in disaggregated lung cells 10 days after SeV challenge (Figure 4C, Supplementary Figure 3A). We also assessed lung injury by hematoxylin and eosin staining of lung tissue 10 days after SeV challenge and found increased inflammation and epithelial cell damage in undepleted mice compared to CD8^+^ T cell-depleted mice (Figure 4D). This supported our hypothesis that CD8^+^ T cells contribute to fatal SeV-induced immunopathology.

**Figure 4.**
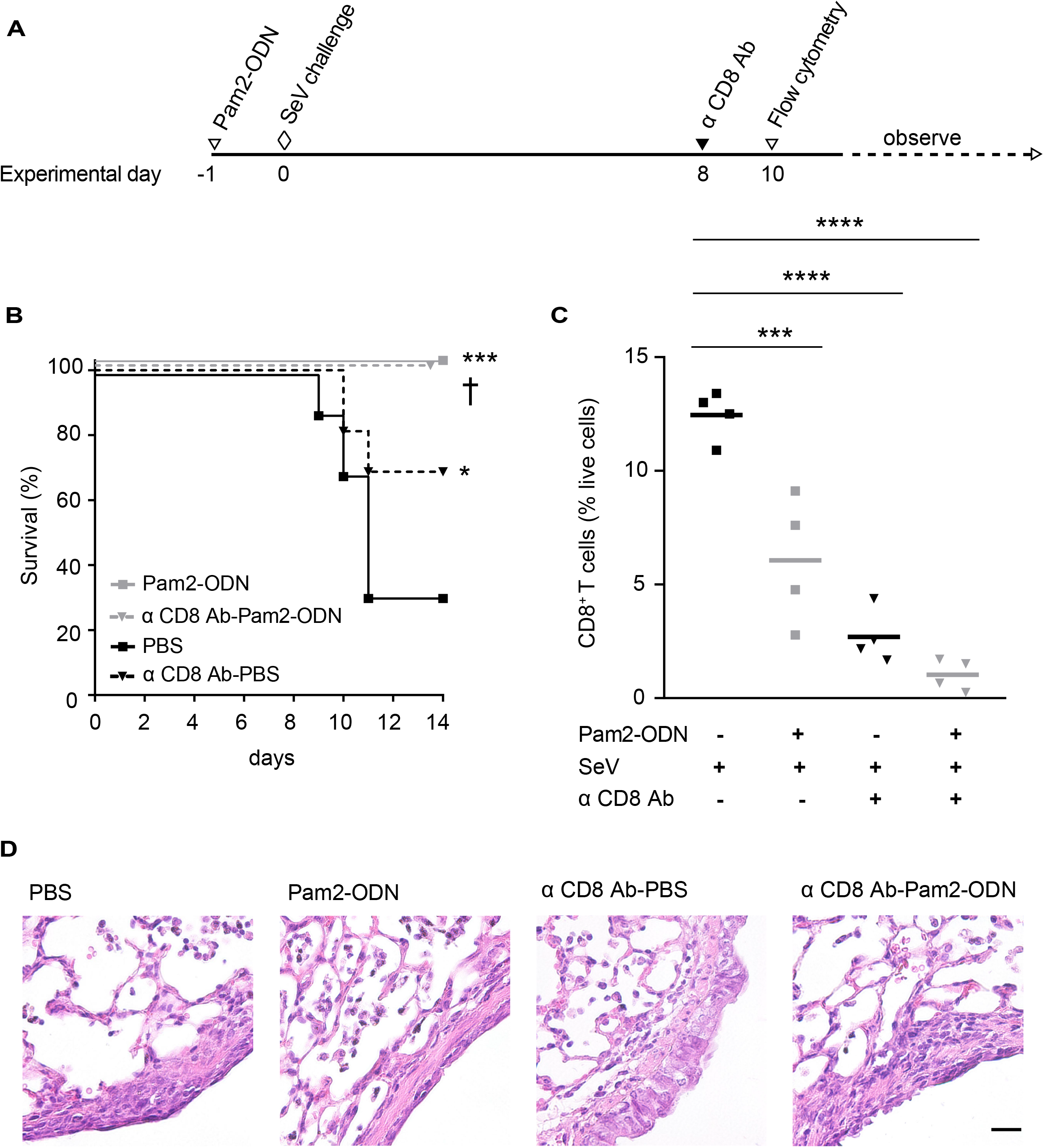
Pam2-ODN treatment reduces CD8^+^ T cell associated SeV induced immunopathology. Experimental outline **(A)**, survival **(B)** and percentage of CD8^+^ T cells **(C)** from disaggregated mouse lungs 10 days after SeV challenge following pretreatment with PBS or Pam2-ODN and with or without CD8^+^ T cells depleted on day 8 of SeV challenge. **(D)** Lung histology 10 days after SeV challenged with or without Pam2-ODN treatment and/or CD8^+^ T cells. Data are representative of two independent experiments. Scale bar = 100 μm. n=16 mice/group for survival in experiment A and n=4 mice/group in experiment B. \*\*\*\**p*<0.0001 compared to PBS in (c), \*\*\**p*<0.0005 compared to PBS in (B) and (C), †*p*<0.05 compared to PBS, \**p*<0.05 compared to PBS.

To assess the role of CD8^+^ T cells throughout the course of infection, mouse CD8^+^ T cells were depleted prior to and during SeV challenge (Figure 4A, Supplementary Figure. 3A, B). This depletion resulted in significantly reduced survival of SeV infection (Supplementary Figure 3C), compatible with the known antiviral functions of CD8^+^ T cells (34–36). However, it is notable that Pam2-ODN treatment still significantly enhanced survival of SeV challenge even in the absence of CD8^+^ T cells (Supplementary Figure 3C). This finding was congruent with our previous studies showing Pam2-ODN inducible resistance against bacterial pneumonia despite the lack of mature lymphocytes (*Rag1*^−/−^) (18).

### Pam2-ODN treatment leads to extracellular inactivation of virus particles

As the antiviral protection consistently correlated with reduced viral burden *in vivo*, and as the reduced virus burden likely contributes to the reduced CD8^+^ T cell levels, we sought to determine how Pam2-ODN-induced responses cause antiviral effects. Assessing the effect of Pam2-ODN on SeV burden in immortalized mouse epithelial cells (MLE-15) and primary mouse tracheal epithelial cells (mTEC), we found that Pam2-ODN treatment reduced SeV burden at every time point measured, reflecting the inducible antiviral capacity of isolated epithelial cells (Supplementary Figure 4). Further, we investigated whether the principal Pam2-ODN effect occurred before (extracellular) or after (intracellular) virus internalization into their epithelial targets. SeV inoculation was carried out at 4° C preventing SeV internalization while allowing SeV attachment to epithelial cells (26–28). Using multiple methods to determine the effect of Pam2-ODN on SeV attachment, we found no differences in attachment (Figure 5A-D). However, even though similar numbers of virus particles were attached to epithelial cells, when these attached virus particles were liberated from the epithelial cell targets, virus particles from Pam2-ODN-treated epithelial cells were less able to subsequently infect other naive epithelial cells (Figure 5E, F). As the number of attached virus particles was the same, this difference in SeV burden in cells that received liberated virus particles from PBS vs Pam2-ODN treated cells indicated that SeV is inactivated prior to epithelial internalization.

**Figure 5.**
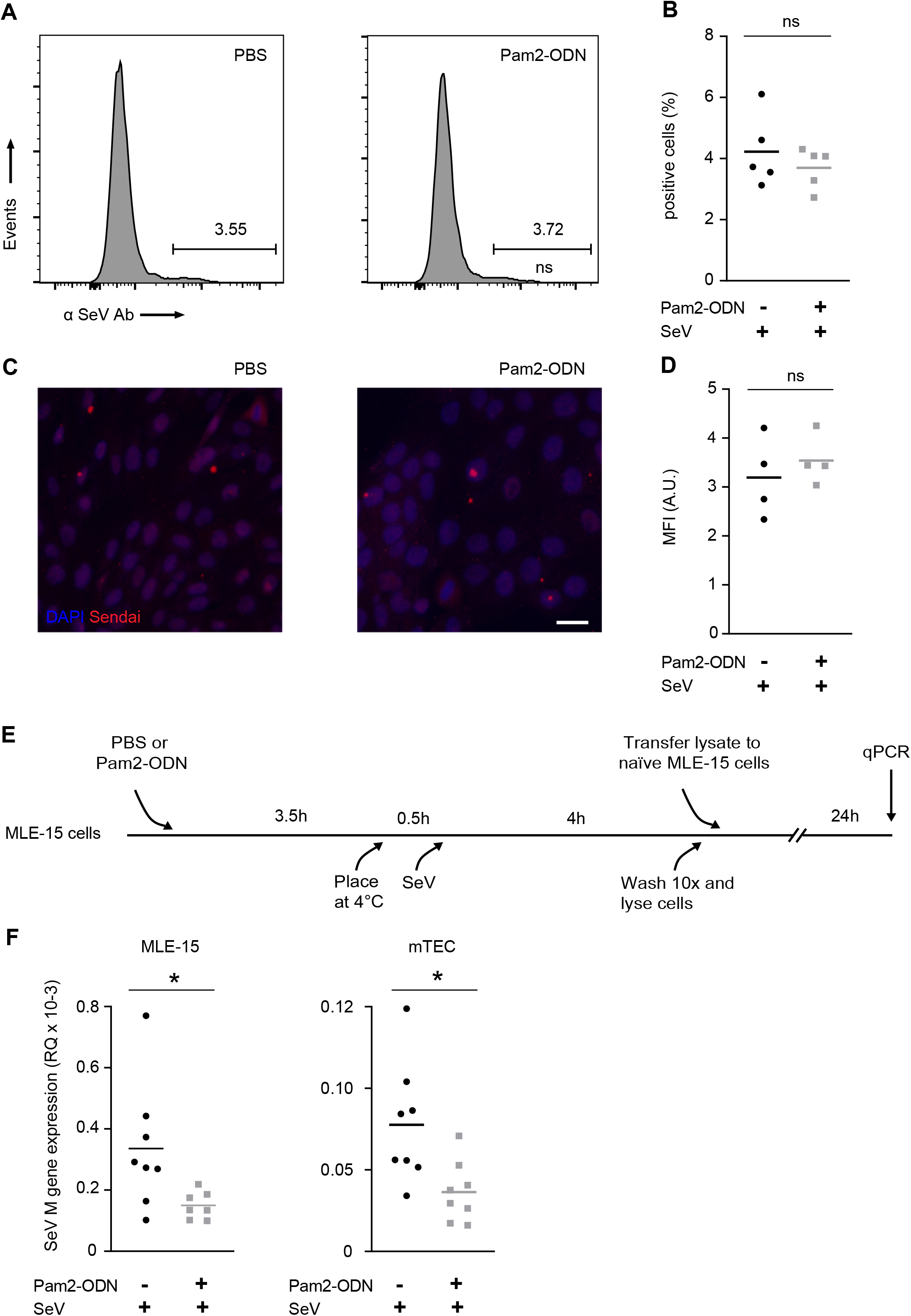
Pam2-ODN inhibits SeV without altering attachment. **(A)** Flow cytometry to measure virus attachment to epithelial cells 4 h after SeV challenge. **(B)** Percentage of SeV positive epithelial cells from **(A)**. **(C)** Representative examples of immunofluorescence for virus attachment. **(D)** Mean fluorescence intensity of SeV-exposed epithelial cells 4 h after SeV challenge. **(E)** Experimental outline showing viral attachment and prevention of virus internalization by epithelial cells. **(F)** SeV M gene expression in untreated MLE-15 cells (left) or primary tracheal epithelial cells (right) challenged with liberated virus (uninternalized virus particles) from cultures that had been pretreated with PBS or Pam2-ODN prior to SeV infection 24 h after transfer of liberated virus to new cells. Data are representative of five independent experiments. \**p*<0.05

### Pam2-ODN-induced epithelial ROS protect against SeV infection and CD8^+^ T cell immunopathology

The anti-influenza response initiated by Pam2-ODN requires epithelial generation of ROS from both NADPH-dependent dual oxidase and mitochondrial sources (22, 23). Extending these findings to the SeV model, an NADPH oxidase inhibitor (GKT 137831) fully abrogated the Pam2-ODN-induced anti-SeV response (Figure 6A). Similarly, treatment with a combination of FCCP (an uncoupler of oxidative phosphorylation) and TTFA (a complex II inhibitor) obviated the Pam2-ODN-induced anti-SeV response (Figure 6B) (22, 23). Further, it was found that Pam2-ODN induced epithelial generation of ROS were required for inactivation of SeV prior to epithelial entry (Figure 6C). Congruent with these *in vitro* and *ex vivo* studies, mice treated with FCCP and TTFA before Pam2-ODN treatment and SeV challenge (Figure 6D) demonstrated reduced survival (Figure 6E), increased SeV burden (Figure 6F), and increased CD8^+^ T cells on day 10 (Figure 6G).

**Figure 6.**
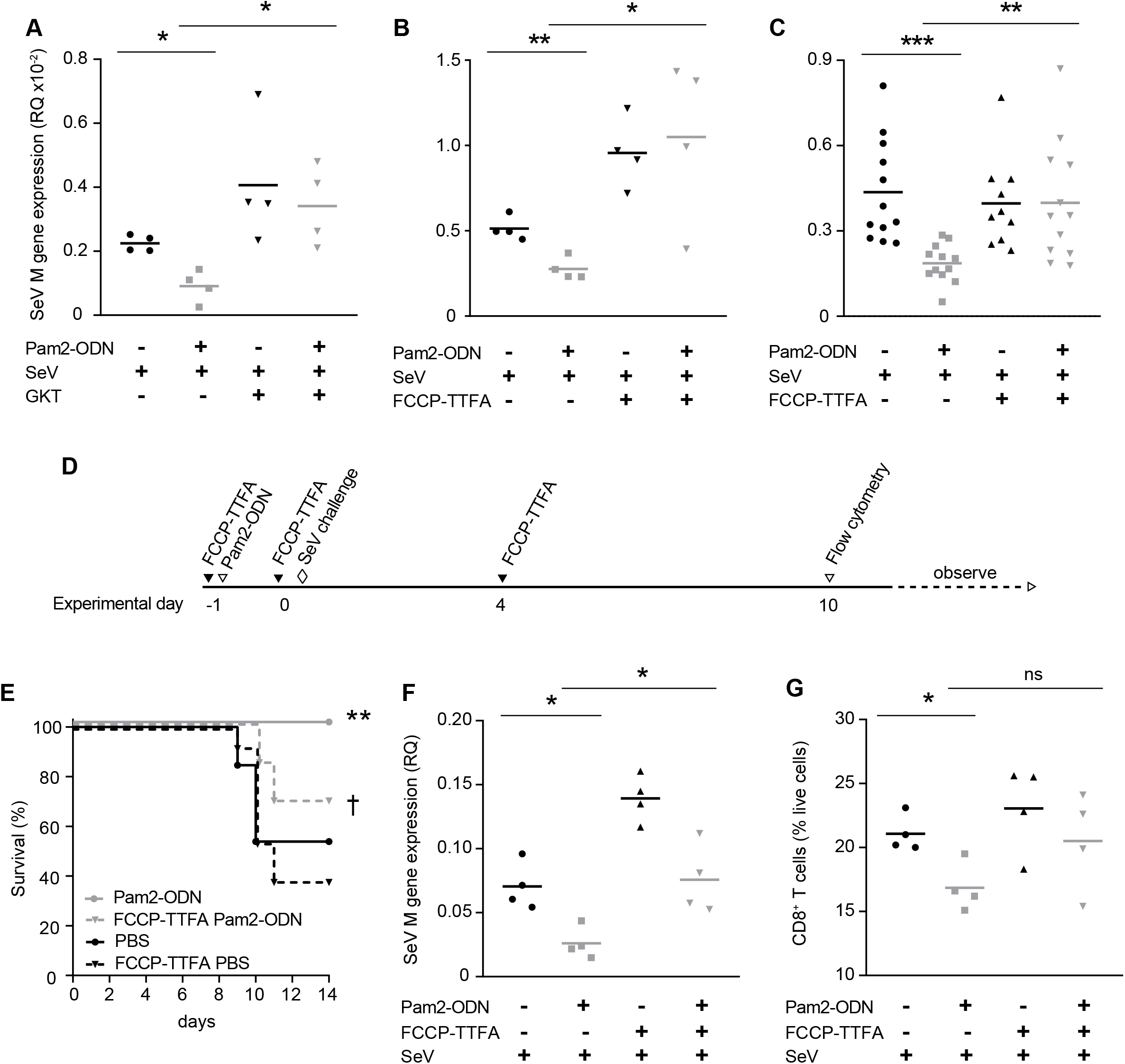
Pam2-ODN induced reactive oxygen species protects against acute SeV virus infections and immunopathology. SeV burden in MLE-15 cells with or without treatment with Pam2-ODN and/or NADPH inhibitors **(A)** or mitoROS inhibitors **(B)**. **(C)** SeV M gene expression in untreated MLE-15 cells challenged with liberated virus from cells that had been pretreated with PBS or Pam2-ODN with or without mitoROS inhibition. **(D)** Experimental outline. **(E)** Survival of SeV challenge in mice treated with PBS or Pam2-ODN and/or mtROS inhibitors. **(F)** Lung SeV burden measured on day 5 and **(G)** lung CD8^+^ T cells assessed on day 10. Data are representative of three independent experiments. n=13 mice/group in experiment **D** and **E**. \*\*\**p*<0.0001, \*\**p*<0.005, \*\**p*<0.01 compared to PBS, †*p*<0.05 compared to Pam2-ODN-treated mice without ROS inhibition, \**p*<0.05.

## Discussion

In this study, we demonstrate that therapeutic stimulation of lung epithelial cells enhances mouse survival of acute SeV infections by both reducing the virus burden and attenuating host immunopathology. While our group has demonstrated inducible resistance against multiple respiratory pathogens including viruses (18–23, 31), these studies demonstrate for the first time when in the virus lifecycle the anti-viral effects begin (viz., prior to internalization), and substantiate the role of ROS in protection against SeV.

While Pam2-ODN treatment provided a significant host survival benefit in SeV infection, we observed this survival benefit occurring after the time when PBS-treated mice had cleared the virus. This observation prompted the hypothesis that host mortality is not the exclusive result of direct viral injury to the lungs, but due at least in part to the host response to the virus infections. We observed enhanced survival of SeV infections in mice depleted of CD8^+^ T cells 8 days after infection (Figure 4A,B), revealing the importance of balancing the dual functions of CD8^+^ T cells in anti-viral immunity and in causing fatal immunopathology. Our findings suggest that the surge in CD8^+^ T cells within the lungs after most virus has been cleared causes physiologic impairment via lung injury and cell death (Figure 4D). These observations demonstrate an advantage of early immune stimulation to enhance viral clearance and late immune suppression to prevent immunopathology and enhance overall outcome of respiratory infections. This is potentially informative in the context of treating pneumonia in immunocompromised patients, and is likely applicable to broader clinical populations, including those suffering lung injury associated with SARS-CoV-2. Based on this reasoning, clinical trials of the use of Pam2-ODN to prevent or treat early COVID-19 have been launched (NCT04313023, NCT04312997), and we suggest that therapeutic targeting of CD8^+^ T cells later in COVID-19 be considered.

Previous reports support the concept of counter-balanced immune protection and immunopathology by CD8^+^ T cells during virus infections (36–41). Some reports have shown that antigen-experienced memory CD8^+^ T cells enhance respiratory syncytial virus (RSV) clearance, but also mediate severe immunopathology (39, 42). However, our study is the first to demonstrate the survival advantage in paramyxovirus respiratory infection of either stimulating the lungs’ mucosal defenses early in the infection or of suppressing the CD8^+^ T cells later in the infection. Our findings are also congruent with reports on the role of CD8^+^ T cells in non-respiratory viral infection models, such as in West Nile virus infection, where CD8^+^ T cell deficient mice display decreased mortality (40). While findings from that study and others reveal that the harmful effects of CD8^+^ T cell mediated immunopathology may supersede the benefits of T cell mediated viral clearance, the question arises of what might be the adaptive value of the vigorous late CD8^+^ T cell response. One possibility is that it might ensure that the infection does not flare again, but that seems implausible since the host has successfully defended itself against the initial infection, and innate immune mechanisms presumably remain intact and are possibly primed (43, 44), in addition to the multiple adaptive immune mechanisms that increasingly come into play. The possibility that the immunopathology simply results from an error on the part of the immune system also seems implausible in view of the substantial rate of host mortality, suggesting there is likely an adaptive value to the response. A third possibility, that the persistence of pockets of low level infection might lead to chronic lung pathology, is supported by a recent study showing that sites of viral RNA remnants following influenza infection are linked to chronic lung disease (45). Thus, a trade-off may exist between the adaptive value of a vigorous CD8^+^ T cell response to prevent chronic lung disease and the acute mortality it can cause. Manipulating this balance therapeutically will need to account for both the benefits and costs of the response. It is particularly appealing to develop inducible anti-microbial strategies that do not rely on conventional T cell-mediated microbial clearance and are also effective in vulnerable immune deficient populations (18, 22, 25, 46).

Although the CD8^+^ T cell depletion studies enhanced our understanding of immunopathology in virus infections, much of the survival benefit against SeV infection was mediated by rapid anti-viral effects induced by Pam2-ODN. This led us to investigate the mechanisms of these inducible anti-viral effects. Given the multiple steps in the virus life cycle, it was not known at what stage Pam2-ODN exerted its anti-viral effect. Exploring this, we found no difference in number of SeV particles attached to the cells between PBS and Pam2-ODN treatment (Figure 5A-D). However, attached virus particles that were liberated from Pam2-ODN treated cells retained less infective capacity when added to naïve epithelial cells, revealing pre-internalization virus inactivation by Pam2-ODN treatment (Figure 5E, F).

Knowing that Pam2-ODN inducible resistance required ROS production to protect against influenza (22), we studied the role of ROS in Pam2-ODN-mediated reduction in SeV burden. ROS inhibition not only led to attenuation of Pam2-ODN’s anti-viral effect but allowed increased lung CD8^+^ T cell numbers, implicating Pam2-ODN-induced ROS in preventing both identified mechanisms of mouse mortality in SeV pneumonia (Figure 6). ROS inhibition led to loss of Pam2-ODN-inducible *in vitro* inactivation of SeV prior to epithelial internalization (Figure 6C), demonstrating for the first time that epithelial ROS directly contribute to virus inactivation.

Production of ROS as a microbicidal mechanism has been widely reported in phagocytic cells (47–49). However, this mechanism has not been extensively studied in non-phagocytic cells (50), where it apparently acts predominantly extracellularly rather than intracellularly as in phagocytes. (Figure 5F, G). These findings of viral inactivation by epithelial ROS production reveal an essential component of inducible epithelial resistance.

Taken together, these findings provide mechanistic insights into the antiviral responses generated by the lung epithelium and the prevention of host immunopathology that may inform future therapeutics to target immunomodulation as a means to improve the survival of respiratory infections in vulnerable populations.

## Acknowledgments and Disclosures

The authors would like to thank Dr. Yongxing Wang for optimization of ROS inhibition experiments. M.J.T., B.F.D., and S.E.E. are authors on U.S. patent 8,883,174, “Stimulation of Innate Resistance of the Lungs to Infection with Synthetic Ligands.” M.J.T., B.F.D., and S.E.E. own stock in Pulmotect, Inc., which holds the commercial options on these patent disclosures. All other authors declare that no conflict of interest exists.

## Notes

1 This study was supported by NIH grants R01 HL117976, DP2 HL123229 and R35 HL144805 to S.E.E.

### Competing Interest Statement

The authors have declared no competing interest.

