## Supplementary Methods for "Immune modulation to improve survival of respiratory virus infections in mice"

**Wali, Supplemental Information**

**Methods**

**Cells**: Mouse lung epithelial (MLE-15) cells were kindly provided by Jeffrey Whitsett, Cincinnati Children’s Hospital Medical Center, and cultured in DMEM with 2% Fetal Bovine Serum (FBS), 1% insulin and transferrin. MLE-15 cells were authenticated by the MD Anderson Characterized Cell Line Core Facility. To harvest tracheal epithelial cells, mice were anesthetized to expose and excise tracheas. These tracheas were then digested in pronase (1.5 mg/ml, Sigma Aldrich) overnight at 4º C. Tracheal epithelial cells were then isolated and cultured on collagen coated transwells in Ham’s F12 media supplemented with differentiation growth factors and hormones as previously described (1, 2).

**TLR treatments and viral challenge:** For *in vitro* treatments, cells were treated with Pam2CSK_4_ (2.2 µM) and ODN M362 (0.55 µM), 4 h before SeV inoculation as previously described (2, 3). For *in vivo* treatments, 10 ml solution of Pam2CSK_4_ (4 µM) and ODN M362 (1 µM) in endotoxin free water was delivered by Aerotech II nebulizer (Biodex Medical Systems) driven by 10 l/min along CO2 (5%) in air for 30 minutes as previously described (2, 3). SeV was purchased from ATCC (Manassas, VA) and grown in Rhesus monkey cells obtained from Cell Pro labs (Golden Valley, MN). For *in vitro* challenges, multiplicity of infection (MOI) = 1 was used. Unless otherwise stated, mice were challenged with 1 x 10^8^ plaque forming units (pfu) in PBS inserted into the oropharynx under isoflurane anesthesia as described (4). Mice were weighed before and daily after challenge as a measure of morbidity and criteria for euthanasia.

**Viral burden quantification**: Viral burden was determined by reverse transcription quantitative PCR (RT-qPCR) of the Sendai Matrix (M) protein normalized to house-keeping gene 18SRNA. For *in vivo* experiments, mouse lungs were collected 5 days after SeV challenge. RNA from mouse lungs was extracted using the Qiagen RNeasy kit. 500 ng of total RNA was converted to cDNA using Biorad iScript cDNA conversion kit. Viral burden was determined by reverse transcription quantitative PCR (RT-qPCR) of the Sendai Matrix (M) protein normalized to house-keeping gene 18SRNA. 18S forward primer – GTAACCCGTTGAACCCCATT; reverse primer – CCATCCAATCGGTAGTAGCG. SeV M gene forward primer –ACTGGGACCCTATCTAAGACAT; reverse primer – TAGTAGCGGAAATCACGAGG. The Limit of quantification (LOQ) was established for the SeV qPCR assay as the highest dilution of the template still maintaining the linearity of the assay. The threshold cycle (C_T_) value of the LOQ was set as the lower limit for the assay.

**Flow cytometry**: For *in vivo* experiments, mouse lungs were perfused with 5 to 10 ml PBS, dissected, cut into 1 mm^3^ pieces, and digested with collagenase/DNAse I (5 mg/ml, Worthington biochemical) for 30 min at 37° C. After digestion, single cells were collected by passing through a 70 µm filter. These single cells were washed with FACS staining buffer (PBS supplemented with 1% FBS) and stained for specific cell types, as indicated in the antibody table (Table 1). For *in vitro* experiments, MLE-15 cells were seeded on 24 well plates for treatment with Pam2-ODN and SeV inoculation. Cells were trypsinized and washed with FACS staining buffer 2X. Cells were blocked in 5% donkey serum for 30 min before proceeding to staining with Rabbit SeV antibody (MBL International) overnight at 4° C, followed by staining with secondary Alexa488 anti-rabbit antibody (Jackson Immunologicals) for 1 h. Cells were fixed and acquired on a BD LSRII (BD Biosciences) for Alexa488 positive cells.

**Epithelial proliferation assays:** Mice were injected intraperitoneally with 0.1 ml EdU (1 mg/mouse). After 24 h, lungs were inflated and fixed with 10% formalin for 24 h at 4° C and then lungs were embedded in paraffin. Paraffin sections were cut into 5 µm transverse sections of the axial airway, between lateral branches 1 and 2. Lung sections were then stained following the Click-iT EdU Imaging Kit protocol for EdU (Abcam) followed by staining with DAPI for 30 min at room temperature. Images were collected using Olympus BX60 microscope using identical parameters for all conditions. Some lung sections were subjected to antigen retrieval and then stained for Ki67 (1:1000; Invitrogen) or cCasp3 (1:500; Cell Signaling). EdU, Ki67 or cCasp3 positive cells were quantified using a cell counter plugin in ImageJ and normalized to DAPI positive cells in every field of view (number of fields surveyed per mouse sample = 3).

**Bronchoalveolar lavage and differential Giemsa staining**: After deep anesthesia, mouse tracheas were exposed, cannulated with a 20-gauge syringe, and instilled with 1.5 ml of PBS. Approximately 1 ml of BAL fluid was collected per sample. The BAL fluid was then spun down at 4º C at 300 *g* to collect the cells in the pellet. The cell pellet was resuspended in 1 ml of ice-cold PBS and 200 μl of this cell suspension was then subjected to cytocentrifugation at 300 *g* for 5 min. Cells were stained with Giemsa stain for differential count determination and total cells were counted by hemocytometer.

**Immunofluorescence microscopy for SeV:** MLE-15 cells were grown on chamber slides (Labtek), treated with Pam2-ODN for 4 h before inoculation with SeV (MOI 1). Cells were then fixed with 2% paraformaldehyde before staining with rabbit anti-SeV antibody (MBL International) and detected using a secondary anti-rabbit antibody. For each experimental condition, specimens were imaged using Olympus BX60 microscope using identical parameters for time of exposure, color intensity, contrast and magnification. Images were then loaded on ImageJ software to calculate mean fluorescence intensity for each group.

**Hematoxylin and eosin** **staining:** Mouse lungs were fixed by intratracheal inflation with 10% formalin for 24 h, and then transferred to 70% ethanol embedded in paraffin. Tissue blocks were then cut into 5 µm sections, mounted onto frosted glass slides, deparaffinized with xylene, washed with ethanol, then rehydrated and stained with hematoxylin and eosin for morphological changes.

**Viral attachment assays:** MLE-15 cells were cultured in 24 well plates or chamber slides for treatment with Pam2-ODN and SeV inoculations. The cells were infected with SeV at 4º C for 4 hours. Thereafter, the cells were vigorously washed 5X with media to remove unattached virus, then harvested to measure uninternalized SeV burden using immunofluorescence or flow cytometry. For RT-qPCR assays, epithelial cells were treated with Pam2-ODN or PBS, followed by SeV infection at 4º C to prevent virus internalization. Virus particles were allowed to attach to the epithelial targets for 4 h at 4º C. These cells were then extensively washed to remove unattached virus particles, and then the cells were lysed by passing through a syringe 10X to liberate the attached uninternalized virus particles. The liberated virus particles were then transferred to naïve epithelial cells that had no prior exposure to Pam2-ODN. SeV M gene expression was assessed by qPCR after 24 h of SeV replication in the new cells. In some experiments, mitoROS inhibitors (FFCP-TTFA) were used before Pam2-ODN treatment to determine the role of Pam2-ODN induced ROS in SeV inactivation prior to epithelial internalization.

**ROS inhibition *in vitro* and *in vivo*:** NADPH oxidase activity was inhibited by exposing the cells to GKT137831 (10 µM; Selleckchem) 12 h prior to treatment with Pam2-ODN or PBS. Mitochondrial ROS production was inhibited using the combination of FCCP (400 nM, Cayman Chemicals) and TTFA (200 µM, Cayman Chemicals) for 1 h before Pam2-ODN or PBS treatment. For *in vivo* experiments, mice were aerosolized with 10 ml TTFA (200 mM) and FCCP (800 µM) 2 h before Pam2-ODN aerosolization and 2 h before SeV challenge and then again 4 days after SeV challenge.

**Statistics:** All statistical analysis was performed using GraphPad Prism software (Version 8 for Windows, GraphPad Software, La Jolla, CA). Data from one representative experiment of at least three independent experiments are presented as mean +/- standard error of biological replicates. To determine pairwise differences in viral burden or cell numbers, Student’s *t* test was used. Mouse survival analysis of viral challenges were analyzed using Mantel-Cox test. One-way analysis of variance (ANOVA) with multiple comparisons was used to determined differences between multiple experimental conditions.
