## Supplementary Figures for "Immune modulation to improve survival of respiratory virus infections in mice"

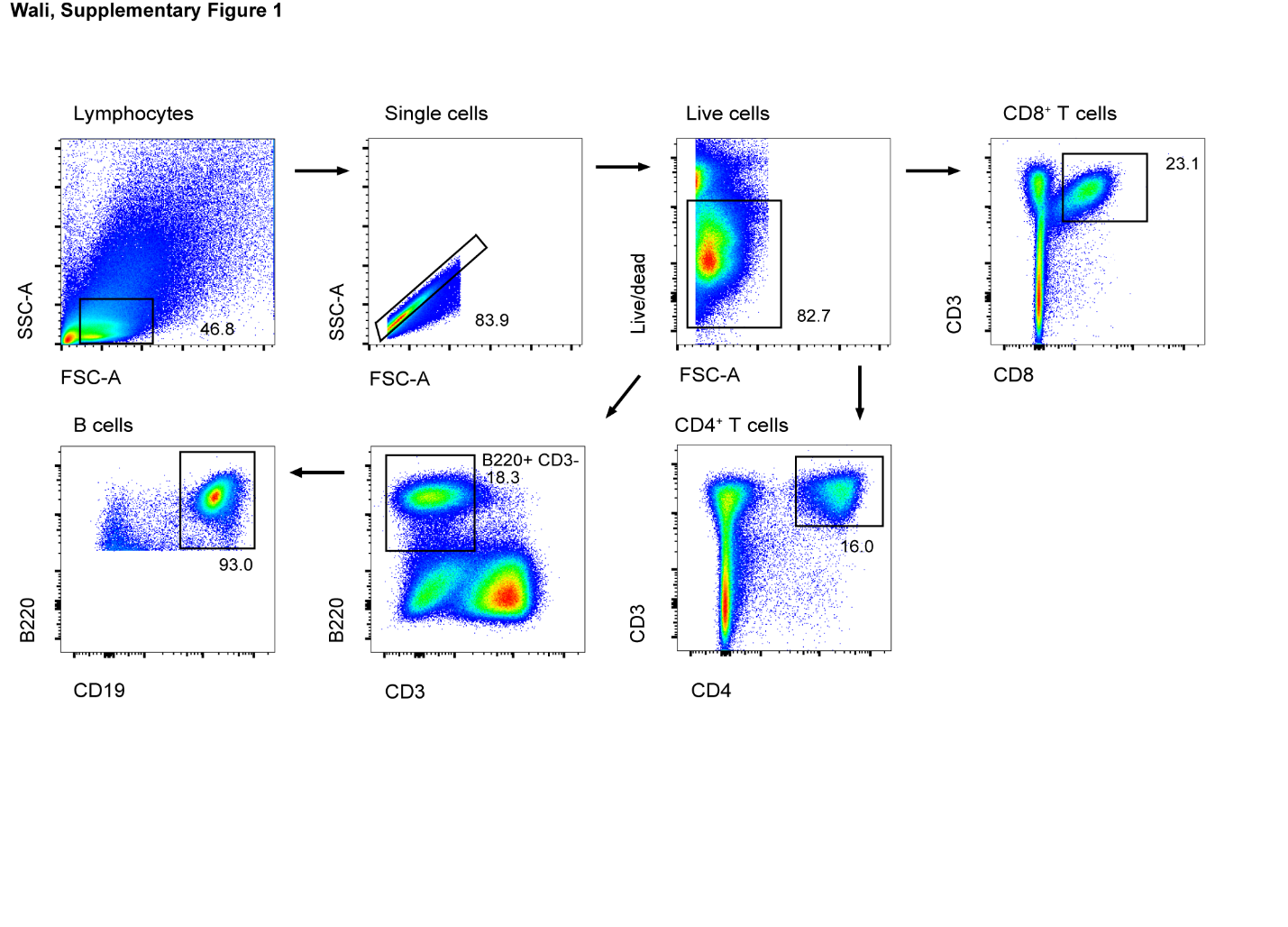

**Supplementary Figure 1.** **Gating strategy for flow cytometry of lung T and B cells.**

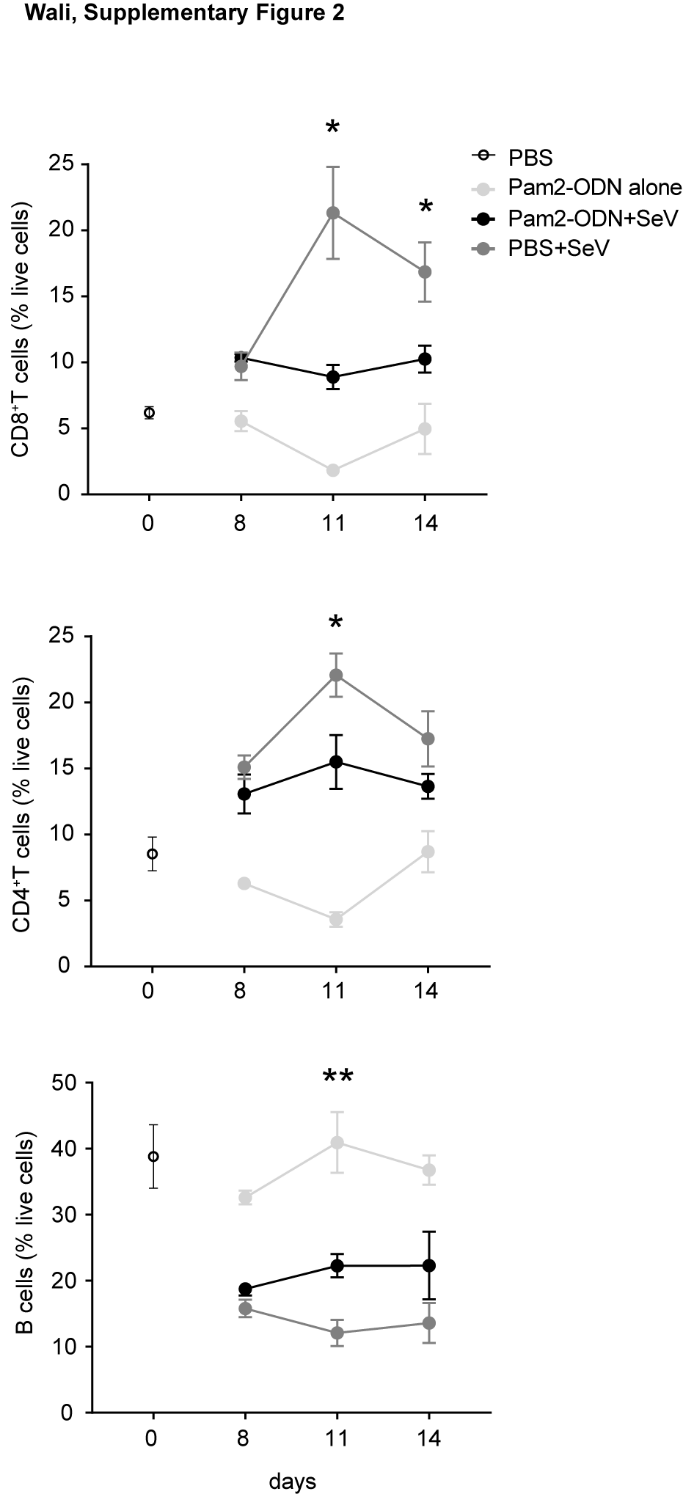

**Supplementary Figure 2.** **Pam2-ODN pretreatment reduced SeV induced lymphocytes.** Disaggregated mouse lung cells positive for CD8^+^ T cells, CD4^+^ T cells, CD19^+^ B220^+^ B cells assessed by flow cytometry in perfused lungs from mice treated with PBS or Pam2-ODN on various days of SeV challenge. Data are representative of three independent experiments. **p*<0.05 compared to PBS+SeV, ***p*<0.01 compared to PBS+SeV.

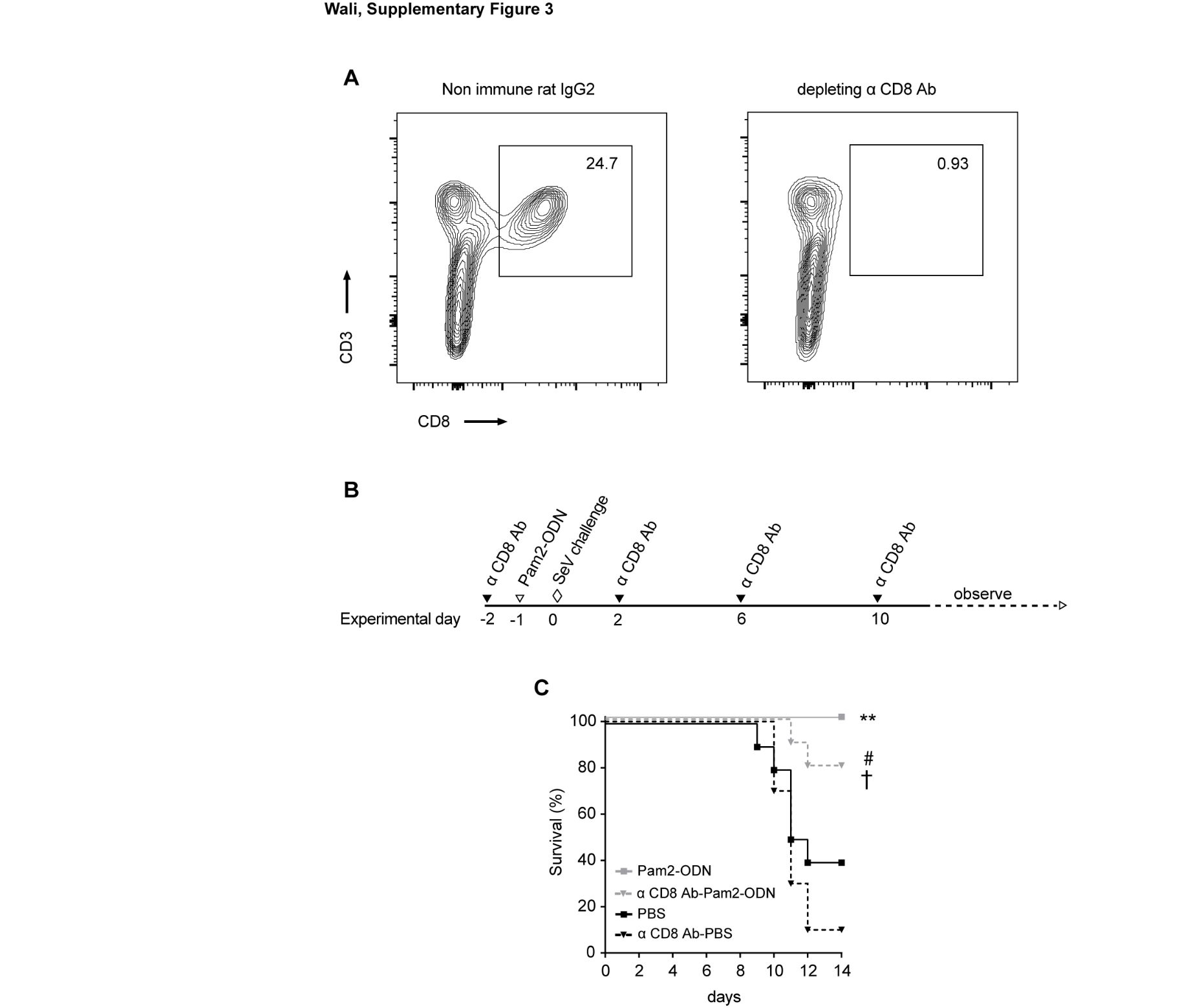

**Supplementary Figure 3. CD8^+^T cell depletion in mice.**  **(A)** Flow cytometry of CD8^+^T cells to confirm depletion of CD8+T cells by depleting antibody. Experimental outline **(B)**, survival **(C)** of mice SeV challenge following PBS or Pam2-ODN treatment and with or without preinfection CD8^+^ T cell depletion. ***p*<0.005 compared to PBS, #*p*<0.005 compared to α CD8 Ab-PBS, †*p<*0.05 compared to PBS.

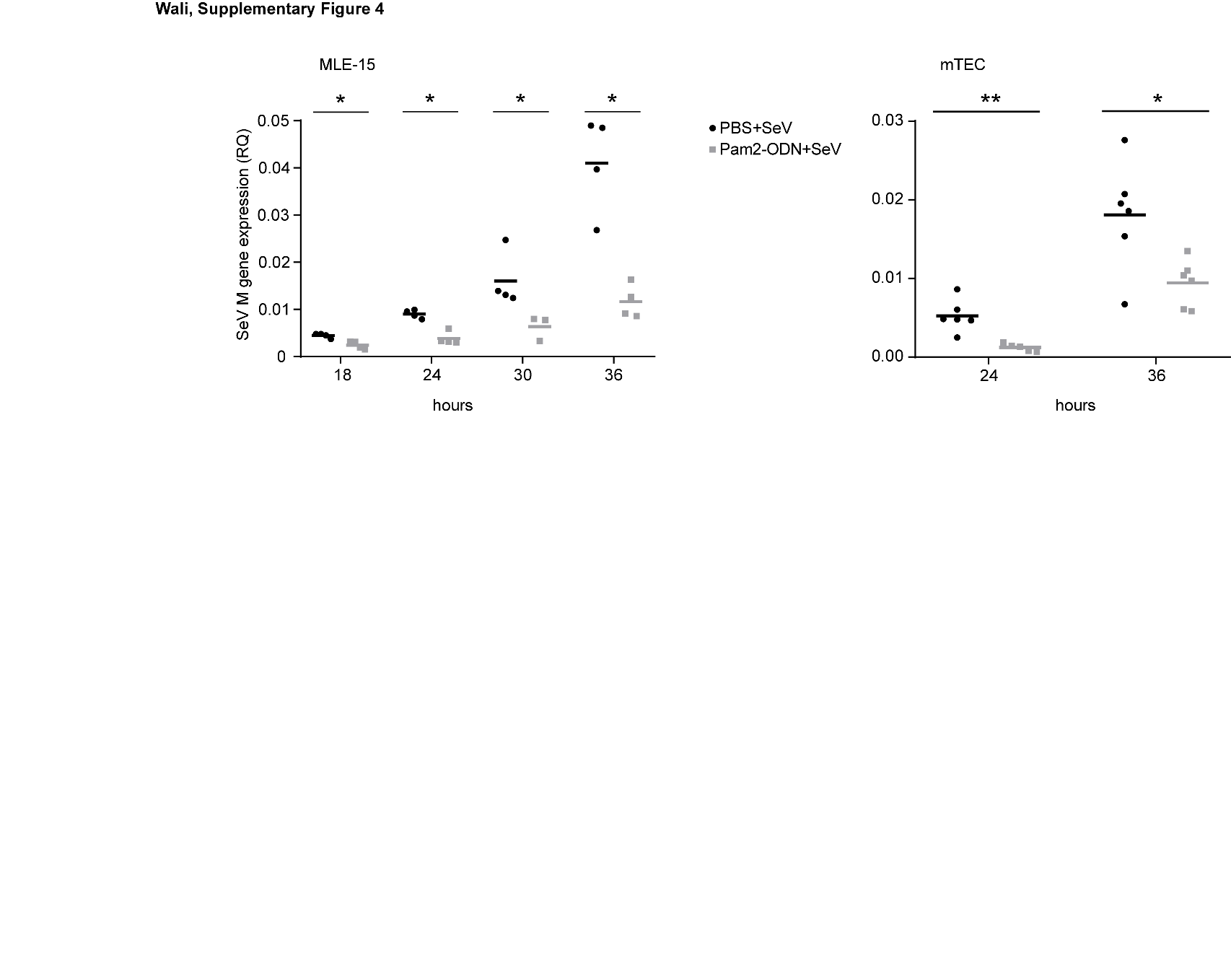

**Supplementary Figure 4**. **Pam2-ODN pretreatment reduced SeV burden in isolated lung epithelial cells.** SeV burden assessed by qPCR in MLE-15 cells and primary mouse tracheal epithelial cells (mTEC) treated with PBS or Pam2-ODN 4h prior to SeV challenge. Data are representative of four independent experiments. **p*<0.05, ***p*<0.005 compared to PBS treated group.
